# Related Topics of a Novel TCR-based Cancer Detection Approach

**DOI:** 10.1101/2020.07.01.182576

**Authors:** Daria Beshnova, Bo Li

## Abstract

We developed a novel algorithm (DeepCAT) to perform de novo detection of cancer associated TCRs, which is based on a convolutional neural network (CNN) model. In this manuscript, we compared its performance with a similar non-deep learning approach, TCRboost, and demonstrated that DeepCAT achieved better prediction accuracy when used to distinguish cancer from non-cancer individuals. Further, although DeepCAT was trained for CDR3s with different lengths, we showed that the combined outcome does not bias the prediction accuracy. Finally, human immune repertoire is affected by many common inflammatory conditions, and our analysis demonstrated that DeepCAT predictions are minimally affected by these factors.

## Introduction

In a previous work, we developed DeepCAT method to perform de novo prediction of cancer associated T cell receptors in the blood TCR repertoire samples (*1*). In brief, DeepCAT is a convolutional neural network (CNN) model that takes the T cell receptor complementarity determining region 3 (CDR3) amino acid sequences as input, and predicts a probability of it is being cancer-associated TCR. True positive cancer associated TCR sequences were derived from tumor infiltrating T cell information (*2*), which was obtained from The Cancer Genome Atlas (TCGA) datasets. Non-cancer TCRs were obtained from the blood TCR repertoire data of healthy donors (*3*). To accommodate CDR3s with different lengths, we trained 5 models each for CDR3s with length 12, 13, 14, 15 and 16. When applied to a TCR repertoire, DeepCAT separately calculates the cancer-associated probability for each TCR, then uses the mean value (also referred as the cancer score) to indicate the cancer status. Contingent on this work, we discussed the issues and topics related to using this deep learning based approach to perform cancer diagnosis. The first section compared the performance of DeepCAT with a non-deep learning based approach; the second section quantified impact of having five different CNN models and the third section discussed how other immunologic conditions may affect cancer score estimations.

## Results

### 1. Comparison of non-Deep Learning approaches and different feature construction methods to predict cancer-associated TCRs

DeepCAT uses principal component analysis (PCA) transformed amino acid feature matrix as input. It is denoted as the “CNN+PCA” approach, as we discuss other potential combinations for the choices of the machine learning method, and the format of input data. Given the differences of amino acid biochemical features (also called amino acid index) between cancer and non-cancer CDR3s, here we applied Adaptive Boosting (*4*) as the non-DL approach to build an ensemble classifier. Below we discuss all possible methods and compare their performances using held-out cancer sample cohorts.

#### i) AdaBoost with 544 biochemical indices

The current amino acid index database (*5*) documented 544 biochemical indices from previous protein structure studies, which can be used as surrogates of the functional and structural impact for amino acids. From the cancer-associated TCRs obtained from TCGA, we removed public TCRs (*6*) that also occurred in healthy individuals, and selected CDR3 sequences with length L between 12 and 16 amino acids (AA) and removed the first 2 and the last 3 AAs without structural contact to the pMHC complex. The total feature set was the union for each informative AA, in other words, the number of features would be (L-5)×544. We used n_L_ to denote the number of CDR3s with length L for cancer CDR3s (derived from TCGA data) and k_L_ for the number for non-cancer CDR3s from VDJdb.

We first subsampled 50% of all the sequences from both populations and used the remaining half of the data for cross validation. For each feature, we compared the 0.5n_L_ cancer observations with the 0.5k_L_ non-cancer ones. If the fold change (cancer over non-cancer) was smaller than 1.1, this feature was removed. Let S denote the number of features left. In the above setting, we have a total of 0.5×(n_L_+k_L_) CDR3 sequences (samples) and S features, with known sample labels (0.5n_L_ with label 1 and 0.5k_L_ with label −1). Let **Y** denote the sample label vector with length 0.5×(n_L_+k_L_) and **X** denote the feature matrix with dimension 0.5×(n_L_+k_L_)-by-S. Based on our analysis of amino acid index differences between cancer and non-cancer TCRs, we determined that the prediction power for individual features is weak. Therefore, we applied Adaptive Boosting algorithm, an ensemble learning approach that is able to aggregate weak classifiers into a stronger one.

Model training was completed using adaboost() function in R package JOUSBoost (*7*), with 50 rounds of boosting and tree depth of 10. We selected parameters based on the criteria of minimizing the number of training cycles (rounds) and the complexity of classification tree (depth) while minimizing cross-validation (CV) errors. CV errors were calculated by applying the trained classifier for CDR3 length L (denoted as T_L_) to the independent validation data with known class labels. We ran the subsampling 10 times and selected the one with the best cross-validation value. The above procedure was repeated for L=12, 13, 15, and 16, but not for L=14, where four-fold cross validation was applied because we found that this setting achieved smaller CV error. Therefore, in total 5 classifiers were trained and were denoted as T_12-16_. We applied TCRboost to define cancer score in the same way as DeepCAT.

#### ii) AdaBoost with PCA encoding

Using the 544-by-20 amino acid index matrix, we performed principal component analysis after removal of missing values (531 features remaining) and selected the top 15 PCs (PC1-PC15, ranked by variance explained). We modified the AdaBoost method by using the 15 PCs to construct ensemble tree classifiers. The final cancer score was defined the same way as DeepCAT.

#### iii) CNN with 544 biochemical indices

We implemented the same CNN network as DeepCAT, with the same number of convolutional layers and the same number of filters for each layer. We encoded each amino acid with 531 features that do not contain any missing values. Therefore, the input image had a dimension of 531-by-L, where L is the length of the CDR3 sequence. The dimensions of the filters of the first CNN layer were 531-by-2. Five models were trained for CDR3s with L=12, 13, …, 16, using the same training data as DeepCAT. Cancer score was calculated as the mean of model predictions for a given TCR repertoire.

#### iv) CNN with PCA encoding

This is the original DeepCAT method previously described. In this model, each CDR3 sequence was first converted into a 2-dimensional image with PCA encoding to integrate the AA indices, and passed onto two consecutive 1-dimensional convolutional layers each with 8 and 16 filters. Random dropouts at a rate of 40% were applied to the dense layer to prevent over-fitting. The output layer generated a probability of cancer association. We built five models each for lengths 12 through 16.

All four models were trained using 20 runs of 4-fold cross validation. To evaluate their performances, we applied each model to the treatment-naïve cancer patient cohorts, including melanoma, breast, ovarian, and pancreatic cancers (*8-11*). We used the 176 age-matched healthy donors from the Emerson et al., 2017 cohort as control (*3*). Prediction power for each method on each cohort was assessed by AUC values (**Figure 1**). Prediction accuracy for CNN+PCA encoding was higher than AdaBoost+PCA encoding. Specifically, by comparing **a**) and **b**), we confirmed that the prediction accuracy for CNN+PCA encoding is higher than AdaBoost+PCA encoding. This result suggests that deep neural network is able to construct non-linear mapping with higher complexity that is superior than traditional approach. To evaluate the value for PCA based feature construction, we compared **a**) with **c**), and **b**) with **d**), and generated the ROC curves for all cohorts combined (**e**). We found that PC encoding only slightly increase (1%) the performance for AdaBoost, but increased the AUC by 6% for CNN. These results suggested that although deep learning was able to construct complex non-linear functions, altering its input by proper feature construction may improve its performance. We believe that this result is informative for the application of deep neural networks to investigate genomics data, because encoding protein and DNA sequences is non-trivial and usually necessary for this type of analysis.

**Figure 1.**
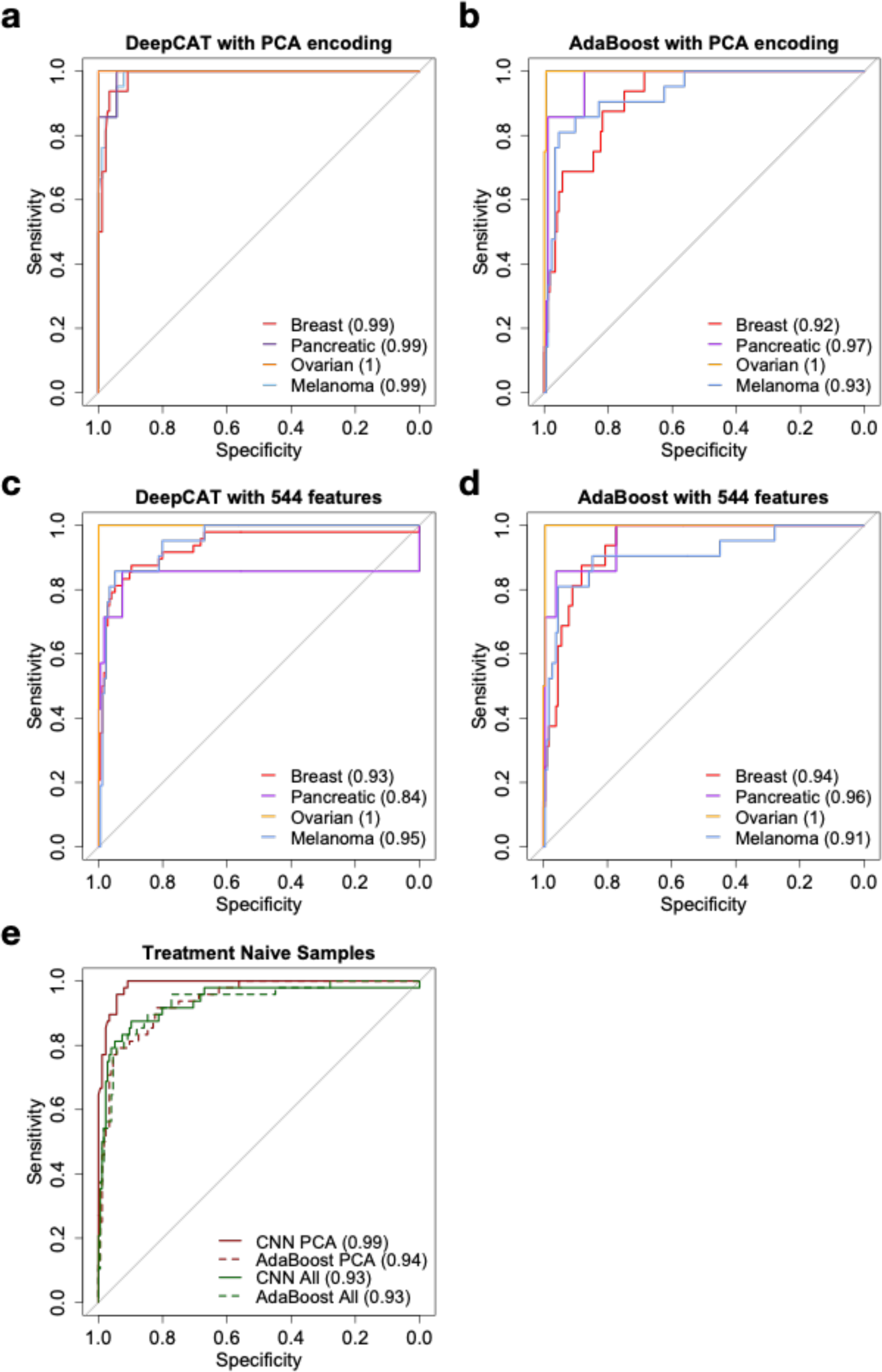
ROC curves for cancer scores predicted using four different approaches. Values in the parenthesis in the legends are AUC values of the ROC curves.

### 2. Distribution of caTCR probabilities from five DeepCAT models and their effects on cancer score estimation

In DeepCAT, we introduced five models each for CDR3s with lengths 12 to 16. The outcome for each model was the probability of cancer association. A CDR3 with higher probability was more likely to be associated with cancer. The distributions of the outcomes from the five models were not identical (**Figure 2**). Specifically, CDR3s with lengths 12, 13, and 16 had similar distributions, whereas lengths 14 or 15 had lower probability. We used the caTCR probabilities predicted from independent testing data.

**Figure 2:**
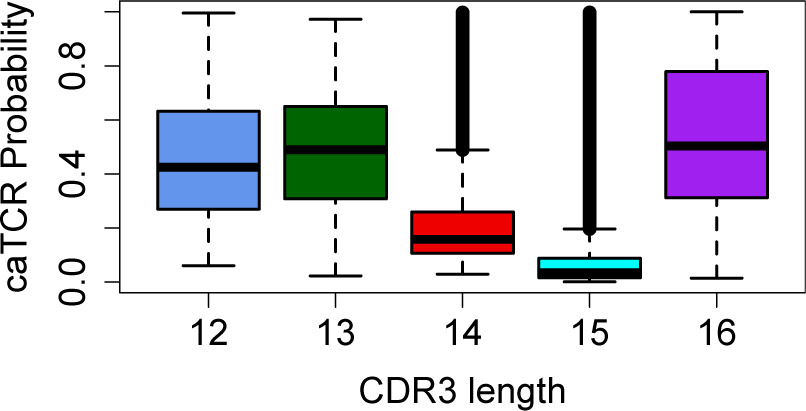
Boxplots showing the distributions of caTCR probabilities estimated from five DeepCAT models.

We next evaluated how this distribution difference affected cancer score calculation and the signals we observed between cancer and normal individuals. We combined all the CDR3s from healthy donor cohorts (Emerson 2017, DeWitt 2015, Kanakry 2016 and Chu 2019) (*3, 12-14*), and from cohorts of patients with cancer. Investigation on CDR3s with lengths 12 to 16 revealed higher usage for CDR3s with lengths 15 and 16 in the cancer cohort compared to normal (**Figure 3**).

**Figure 3:**
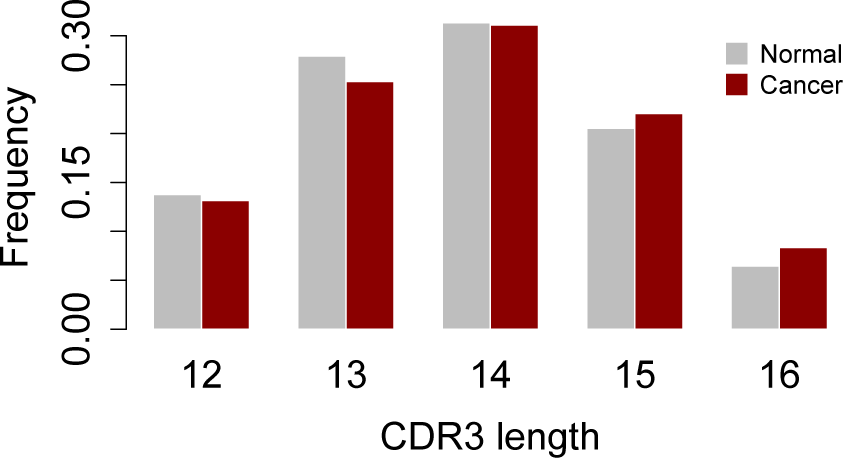
Barplots showing the frequency distribution of CDR3 lengths in normal or cancer cohorts.

We implemented in silico simulations to test how these differences affect cancer score estimation under the null hypothesis that there is no difference in caTCR probability distribution between individuals with and without cancer. Specifically, we simulated 100 ‘patients with cancer’ and 100 ‘normal individuals’. Each individual had 500 TCRs, with lengths following the distributions of caTCR probabilities (**Figure 3**) and with different CDR3 lengths for cancer or normal. The numbers for each CDR3 length were sampled using Multinomial sampler in For each individual, we sampled the number of length L CDR3s following the caTCR length distribution of the healthy donor (**Figure 2**). For example, if individual with cancer #1 has 72 sequences with length 16, we sampled 72 numbers from the caTCR probabilities estimated from length 16 CDR3s. We used the same caTCR probabilities for both individuals with cancer and healthy controls under the null hypothesis. We did not observe higher cancer scores for the patients with cancer in this analysis (**Figure 4**). In other words, adjusting for five DeepCAT model outcomes and CDR3 lengths, the expected null distributions of cancer scores for individuals with or without cancer were similar. This result indicated that the observation of higher cancer scores in patients with cancer in our study was not an artifact of different CDR3 lengths or caTCR probability distributions.

**Figure 4.**
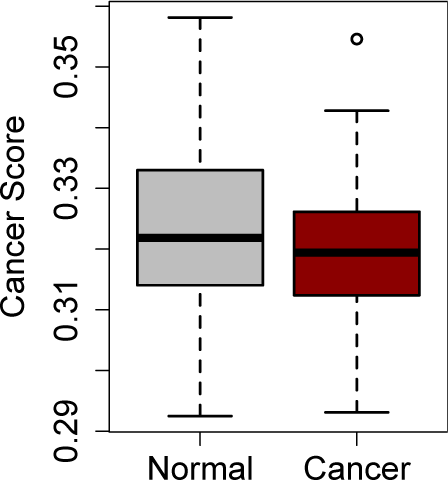
Cancer scores estimations from simulated 100 cancer and 100 normal individuals under the hypothesis that caTCR probabilities follow the same distribution in cancer and normal cohorts.

### 3. Influence of non-cancer chronic inflammatory conditions on the cancer score

Chronic inflammation is common among the population, and it includes chronic viral infection, auto-immune disorders, and cancer. In this work we have demonstrated increased cancer scores in patients with malignant tumors, but it remains unclear how non-cancer-related chronic inflammation may affect the cancer score. To investigate, we collected 3 cohorts, including those with HCMV infection (*3*) (Emerson 2017), rheumatoid arthritis (RA) (*15*), and multiple sclerosis (MS) (*16*). An advantage of these cohorts was that they had healthy donor samples uniformly profiled with the patient samples. However, unlike for Emerson 2017 (*3*), the other two cohorts could not be compared to other samples in our analysis, because the Savola cohort (*15*) used flow-sorted CD8+ T cells, and the Alves Sousa cohort (*16*) was profiled using 5’ RACE with mRNA. For all three cohorts, cancer scores were increased in patients with inflammatory conditions (**Figure 5**), but this increase (ratio of means) did not reach the magnitude seen in the cancer patients.

**Figure 5:**
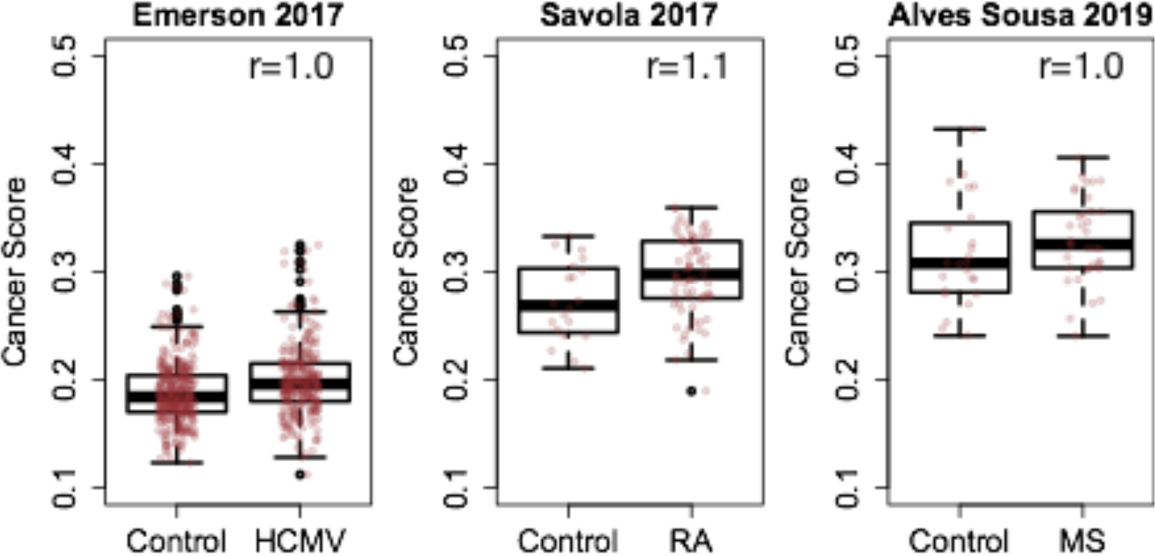
Non-cancer chronic inflammatory conditions might slightly raise cancer scores. r is the ratio of means between the diseased groups over control.

In conclusion, pre-existing chronic inflammatory conditions slightly increase cancer scores, which may result in a reduction in diagnostic specificity when the cancer score is applied to the general population. This caveat, however, can be potentially addressed by exhaustive examination of each patient’s medical history for chronic viral infections and common autoimmune disorders. Future efforts will be needed to generate more uniformly profiled TCR-seq data for other chronic conditions and explore how they affect cancer scores. With enough data, better Deep Learning models can be developed to differentiate patients with cancer from individuals without cancer but with other inflammatory conditions.

## Acknowledgements

We thank the reviewers of the DeepCAT manuscript *(1)* for bringing up with the issues discussed in this manuscript and thoroughly reviewing the results. This work is supported by CPRIT RR170079 (BL).

## Data and Code Availability

Source codes of TCRboost and DeepCAT are available at: https://github.com/s175573/DeepCAT.

